# Zdhhc17/Hip14 functions in neurons to control behavior via DHHC-dependent and independent mechanisms

**DOI:** 10.64898/2026.11.12.766291

**Authors:** Yuzhu Cheng, Nicholas J. Santistevan, Kade Fink, Seth Jacobson, Christian Mosimann, Alexa Burger, Jessica C. Nelson

## Abstract

To navigate a complex sensory environment, animals filter out inconsequential sensory inputs and habituate to familiar stimuli. By establishing innate response thresholds and modifying them through habituation, animals can focus on key sensory stimuli while ignoring irrelevant ones. Huntingtin-interacting protein 14 (Hip14), also known as zDHHC17, is a palmitoyl acyltransferase (PAT) that regulates both of these sensory filtering processes. Larval zebrafish lacking Hip14 are hypersensitive to acoustic stimuli and cannot regulate their responses to repeated acoustic stimuli via habituation. Yet how Hip14 regulates these behaviors, and whether it does so through different mechanisms, is unknown. Prior work demonstrates that Hip14 is expressed throughout the nervous system, but where it functions to control these overlapping sensory functions remains unaddressed. We found that the catalytic (DHHC) domain, despite being a defining feature for PATs including Hip14, is not required for regulating acoustic startle sensitivity, but is sufficient to regulate habituation. We find, furthermore, that Hip14 function in either mechanosensory hair cells or neurons can restore normal startle sensitivity, whereas habituation is regulated exclusively by neuronal expression. Finally, we found that the Hip14 N-terminal ankyrin repeat domains are crucial for its control of behavior. Together, our study establishes that habituation and startle sensitivity are regulated through distinct circuit and molecular mechanisms and establishes a new in vivo catalytic-independent role for Hip14.

**Significance Statement:** Sensory dysfunction is a common feature of many neurological disorders. Hypersensitivity to stimuli commonly co-occurs with deficits in the ability to habituate to familiar stimuli, yet how these processes are related is not understood. Using larval zebrafish, we found that a neuronally expressed palmitoyl acyltransferase (PAT) enzyme, Hip14, controls acoustic startle sensitivity through a mechanism that does not depend on its catalytic domain. Conversely, Hip14 regulates habituation through a mechanism that relies on this key domain. Our findings underscore the close connection, but genetic separability of the mechanisms underlying sensory threshold establishment and habituation-mediated threshold modification. Furthermore, our study adds to a growing appreciation for catalytic-independent functions of PATs and lays the foundation for understanding how these enzymes control behavior.

## Introduction

To filter extraneous sensory stimuli, organisms establish innate response thresholds during development and later can modulate them through simple plasticity mechanisms such as habituation (Takahashi et al., 2002, 2017; Burgess and Granato, 2007a; Rankin et al., 2009; Roberts et al., 2011; Wolman et al., 2011; McDiarmid et al., 2017, 2019). Disrupted sensory filtering and habituation are linked to neurological disorders including anxiety (Bakker et al., 2009), autism spectrum disorders (Schoen et al., 2008; Guiraud et al., 2011; Chamberlain et al., 2013; Takahashi et al., 2016), attention-deficit/hyperactivity disorder (Jansiewicz et al., 2004; Panagiotidi et al., 2018), and schizophrenia (Geyer et al., 1990; Braff et al., 1992). Despite their importance to human health, how innate sensory thresholds are established and modulated, including through habituation, remains unclear.

To better understand how animals establish and dynamically modulate behavioral thresholds, we examined a simple behavior, the acoustic startle response, in larval zebrafish (Kimmel et al., 1974; Burgess and Granato, 2007a; Wolman et al., 2011). Larval zebrafish establish innate response thresholds for acoustic/vibrational stimuli; stimuli above the threshold elicit a startle response, while stimuli below the threshold are ignored (Kimmel et al., 1974; Burgess and Granato, 2007a; Marsden and Granato, 2015; Takahashi et al., 2017). Furthermore, zebrafish can also learn to ignore repeated stimuli through habituation at 5 days post-fertilization (dpf) (Roberts et al., 2011; Wolman et al., 2011). Using this system, we previously identified the palmitoyl acyltransferase Huntingtin-interacting protein 14 (Hip14 / zDHHC17) as a regulator of innate acoustic startle sensitivity and habituation (Wolman et al., 2015; Nelson et al., 2020). Loss-of-function mutations in *hip14* lead to hypersensitivity to acoustic stimuli and defects in startle habituation. Moreover, heat-shock-induced ubiquitous expression of *hip14* rescued both phenotypes within hours, suggesting that Hip14 regulates circuit function rather than development (Nelson et al., 2020).

Across vertebrates, acoustic stimuli are detected by mechanosensory hair cells in the inner ear (Whitfield et al., 2002; Schwander et al., 2010). VIIIth cranial nerve afferents transmit sound information to hindbrain neurons that mediate the acoustic startle response. In the larval zebrafish, the acoustic startle response is characterized by a short-latency “C-bend” mediated by reticulospinal Mauthner cells (M-cells) (Kimmel et al., 1980; Korn and Faber, 2005; Burgess and Granato, 2007a). During habituation, the lateral dendrite of the M-cell undergoes synaptic depression (Marsden and Granato, 2015), whereas in Hip14 mutants, the calcium signal within the lateral dendrite remains unaltered after repetitive stimulation (Nelson et al., 2020), indicating a role for Hip14 in regulating activity within components upstream of, or within, the M-cell. Here, we examined Hip14 expression within the M-cell, VIIIth nerve, and mechanosensory hair cells and dissected its role within this circuit.

Beyond its localization, we sought to understand the molecular mechanisms through which Hip14 controls sensitivity and habituation of the acoustic startle response. As a palmitoyl acyltransferase, Hip14 relies on its DHHC (asp-his-his-cys) domain to catalyze palmitoylation of target substrates. Yet, Hip14 has molecular functions independent of the DHHC domain. For instance, *in vitro,* Hip14 facilitates JNK3 phosphorylation through a function that does not rely on the DHHC domain (Yang and Cynader, 2011). Although most PATs contain a DHHC domain, they are otherwise structurally divergent. Specifically, Hip14 and the closely related Hip14L contain N-terminal ankyrin repeats (ARs) not found in other vertebrate PATs. These ARs facilitate recognition of palmitoylation substrates, including a handful that are localized to synapses, for example cysteine-string protein (CSP) and synaptosomal-associated protein 25 (SNAP25) (Lemonidis et al., 2014, 2015, 2017; Verardi et al., 2017). Quantitative assays of sensory-evoked behaviors in the larval zebrafish provide a powerful system for determining the *in vivo* systems-level requirement for the AR and DHHC domains.

Here, we establish the *in vivo* tissue and subcellular localization of endogenous Hip14 within the acoustic startle circuit. Next, we used genetic and molecular techniques to elucidate the contributions of both the DHHC and AR domains to controlling behavior. We uncover parallel contributions of Hip14 within both peripheral sensory and neuronal cells to regulate acoustic startle thresholds. Moreover, we present evidence that Hip14 regulates startle response thresholds through a DHHC-independent mechanism, whereas acoustic startle habituation requires the catalytic domain. Our findings provide evidence for the *in vivo* behavioral relevance of palmitoylation-independent functions for Hip14.

## Materials and Methods

### Zebrafish

All zebrafish use and procedures were approved by the University of Colorado Anschutz Medical Campus Institutional Animal Care and Use Committee (IACUC). All experiments were conducted in the TLF strain. Adult zebrafish were raised on a 14h:10h light-dark cycle. Zebrafish larvae were raised at 29°C in E3 media at appropriate densities: 45–60 embryos per 10 cm dish. Dead or deformed animals were removed daily, and E3 media was changed at 0 and 3 days post-fertilization (dpf). Since sex is indeterminate until later stages, this study did not differentiate larval or embryonic zebrafish by sex.

### Transgenic constructs and mutants

*hip14^p174^* mutants were uncovered in a previous genetic screen (Wolman et al., 2015). All transgenes used for behavioral rescue were inserted into the *pIGLET14a* locus using phiC31 integrase-mediated transgenesis (Lalonde et al., 2024), denoted as *p14a*. To generate *p14a.Huc:hip14-egfp* and *p14a.myo6b:hip14-egfp*, we conducted Gateway cloning, combining *pME-hip14-egfp* with *p5E-myo6b* (Kindt et al., 2012) or *p5e-elav3l* (HuC) (Addgene: #72640), *pDestattB* (pCM268, Addgene# 68313) (Lalonde et al., 2024), and Tol2kit *#302 p3e-polyA* (Kwan et al., 2007). To generate *p14a.Huc:hip14-N90AW118A-egfp* and *p14a.Huc:hip14-AR-deletion-egfp*, we employed site-directed mutagenesis to modify the *pME-hip14-egfp* plasmid either by mutating Asparagine 90 and Tryptophan 118 to Alanine or by excising the 77-249 bp AR region.

### neuroD

*sypb-mScarlett* was graciously gifted by Bruce Appel (Hughes and Appel, 2019). M-cells were labeled by crossing *Tg(gffdmc130a)* (Pujol-Martí et al., 2012) and *Tg(UAS:gap43-citrine)* (Lakhina et al., 2012), respectively.

To generate the *hip14-V5* knock-in, we employed the CRISPR-Cas9 knock-in technique and V5 epitope tagging approach as described by Burg and colleagues (Burg et al., 2016). For oligo sequences, see **Supplemental Table S1**.

For generation of *p24b.ubi tdTomato-GM130* (in *pIGLET* landing site *pIGLET24b*)(Lalonde et al., 2024), we amplified *tdTomato-GM130* from a plasmid generously provided by the Solnica-Krezel lab (Sepich and Solnica-Krezel, 2016), adding restriction sites for subsequent cloning. Next, we ligated the sequence into an existing *pME* vector (**Supplemental Table S1**). The final Gateway construct was generated combining *pENTR5’_ubi* (pCM206)(Addgene#27320) (Mosimann et al., 2011), *pDESTattB (pCM268)* (Mosimann et al., 2013)(Addgene# 68313), and Tol2 Kit #302 (Kwan et al., 2007) using LR Clonase II Plus and Multisite Gateway-optimized vector ratios (Kwan et al., 2007; Mosimann, 2022).

Injection mixes consisted of 25 ng/µL plasmid, 12.5 ng/µL integrase, and 1:10 phenol red (Fisher # 25300054), brought to 10 µL with DEPC water. Injection mixes were injected into offspring at the one-cell stage, arising from crosses between *pIGLET14a* heterozygous carriers and *WT* (*TLF*) adults, carrying at least one copy of the *pIGLET14a* locus (Lalonde et al., 2024). *p24b.ubi:tdTomato-GM130* was generated by injection into embryos from an incross of *pIGLET24b* landing site carriers. Then we screened and selected single transgenic carriers for positive germline transmission, outcrossed transgene-positive F1 animals into the *TLF* background, and ultimately into the *hip14^p174^* heterozygous background. When possible, we identified transgene-carrying animals via expression of visible fluorescent markers between 3-6 dpf using a fluorescent stereomicroscope (Leica M205FCA), and/or genotyped them as previously described (Lalonde et al., 2024)(**Supplement Table S1**). All animals used for behavior were offspring of at least the F2 generation.

### Behavioral Testing

Animals to be tested were first acclimated for at least 30 minutes in the behavior room within a temperature range of 26-29 °C. We recorded all experiments with a high-speed camera (FASTCAM Mini UX50 Type 160K-M-32G) placed above the testing plate. An LED light was positioned above the plate to provide white light illumination. To test for acoustic startle behaviors, 5-6 dpf zebrafish larvae were loaded onto a custom-made acrylic 6 × 6 well plate attached to a mini-shaker (Brüel & Kjær, Model 4810). Acoustic stimuli at six intensities (0.537*g*, 1.279*g*, 2.559*g*, 5.110*g*, 15.350*g*, 51.177*g*) were administered and calibrated with an accelerometer (PCB Piezotronics, Y355B03). For assays of acoustic startle sensitivity, 5 stimuli were repeated at each intensity level with an inter-stimulus interval (ISI) of 40 seconds. The test for acoustic startle habituation ensued, using the highest intensity stimulus (51.177 *g*), repeated 40 times with an ISI of 3 seconds. To assess dark flash responses, 6 dpf larvae were first acclimated on the testing plate to white light conditions for 15 minutes. Next, lights were extinguished 5 times with an ISI of 60s. After behavior testing, video files were background-subtracted using ImageJ/FIJI, and subsequently, larval movements were tracked and analyzed using FLOTE and BATCHAN software (Burgess and Granato, 2007b; Schloss et al., 2025).

### Genotyping

Individual animals were loaded into 96-well plates, methanol-fixed, and then gDNA was extracted in 20 µL of Base solution (0.025 M NaOH, 4 mL 0.2 mM EDTA), heated at 95°C for 15 minutes, then neutralized with 20 µL 0.4 M Tris-HCl pH 5.0 solution. *hip14^p174^* was genotyped using standard PCR techniques with OneTaq (NEB #M0509L) or Q5 High-Fidelity DNA Polymerase (NEB #M0491) followed by a 6-hour restriction enzyme digest with EcoRV-HF (NEB#R3195S). For all primers/oligos used in cloning and genotyping, see **Supplemental Table S1.**

### Statistical Methods

After video tracking and behavior calling using FLOTE and Batchan, we used Microsoft Excel to compile raw data. Wells for which tracking was unsuccessful for >40% of stimuli were removed from analysis. Genotypes were then manually entered into Excel sheets for each animal. Statistical analysis used the R package glmmTMB to fit a logistic mixed-effects model for acoustic startle sensitivity assays. Average responses across animals at each stimulus intensity were then graphed using GraphPad Prism. For acoustic startle habituation, habituation % was calculated as previously described: % Habituation = (1-[response frequency Stimuli 31-40]/[response frequency baseline])*100 (Wolman et al., 2011; Nelson et al., 2020) and graphed using GraphPad Prism. To assess statistical significance for the habituation assay, we performed two-way ANOVA with Tukey’s post-hoc test. For visually evoked behaviors, responses were averaged, graphed, and analyzed using GraphPad Prism, two-way ANOVA, and Tukey’s post-hoc test. Code used in this study is attached as PDF files in the supplemental files.

### Immunohistochemistry and Confocal Microscopy

For immunohistochemistry (IHC), we fixed 5-6 dpf zebrafish using 2% trichloroacetic acid in 1xPBS for 3 hours at room temperature, then washed 5x for 5 minutes in 0.5% TritonX in 1xPBS (1xPBST), blocked with 2% Normal Goat Serum, 1% BSA, and 1% DMSO, in 1x PBST (0.25% Triton X) at room temperature for 1 hour as previously described (Martin et al., 2022). Thereafter, samples were incubated in primary antibodies: Rabbit anti-CX36 Polyclonal 1:500 (Invitrogen #3214145), Mouse anti-V5 IgG2a 1:350 (Fisher #R96025), Chicken anti-GFP 1:500 (Antibodies Inc #GFP-1010), Rabbit anti-DsRed 1:200 (Takara #632496), Rabbit polyclonal anti-Vglut3 1:1000 (Obholzer et al., 2008). After washing in 1x PBST (0.5% Triton X) 5x for 15 minutes at room temperature, secondary antibodies were used at 1:500, including Goat anti-Rabbit Alexa 546 (Fisher #A11035), Goat anti-Mouse IgG2a Alexa 647 (Fisher #A21241), and Donkey anti-Chicken 488 (Fisher #NC0456003), accordingly.

For *neuroD-sypB-mScarlett; hip14-V5* staining, we raised larvae in Phenylthiourea solution (0.03 mg/ml 1x PTU) in E3 media to prevent development of epidermal melanin. PTU solution was changed every day. We fixed 5 dpf larvae using 4% paraformaldehyde in 1xPBS for 1 hour at room temperature then overnight at 4°C. After 3 washes in PBST, we performed antigen retrieval with 150mM Tris-HCl pH 9 for 5 minutes at room temperature, followed by 15 minutes at 70°C. After 2 fast rinses and a 5 minute wash in 1×PBST (0.25% TritonX), samples were permeabilized in 1ml 0.05% Trypsin-EDTA for exactly 40 minutes as previously described (Randlett et al., 2015; Nelson et al., 2023). Samples were then rinsed 2 times in 1×PBST (0.25% TritonX), washed for 10 minutes, and blocked at room temperature for 1 hour with 2% NGS solution as described above. Primary antibodies used were Mouse anti-V5 IgG2a 1:500 and Rabbit anti-DsRed 1:200, followed by Goat anti-Mouse IgG2a Alexa 488 (Fisher # A21131) and Goat anti-Rabbit Alexa 546, both at 1:500.

Post-IHC samples were washed 5× for 5 minutes in 1× PBST (0.25% Triton X), then subjected to stepwise dehydration in 25%, 50%, and 75% glycerol in 1x PBS. Samples were stored in Vectashield (Fisher # NC9265087) at 4°C until use. All antibody solutions were made with 1%BSA, 1%DMSO, in 1xPBST (0.25% Triton X). Prior to imaging, we fully dissected larval brains or removed jaws to visualize hair cells from the ventral side. Images were acquired using a spinning-disk confocal microscope with a 40x objective or 63x objective. Each IHC experiment utilized at least 8 zebrafish samples for each genotype. We performed image processing, including brightness and channel adjustments, in ImageJ.

## Results

### Hip14 regulates acoustic and visual startle sensitivity

To examine sensory-evoked behaviors in larval zebrafish, we used well-established assays for acoustic startle sensitivity and acoustic startle response habituation (Burgess and Granato, 2007a; Wolman et al., 2011; Marsden et al., 2018; Ortiz et al., 2024; Schloss et al., 2025) **(Fig.1A-D)**. As we showed previously (Wolman et al., 2015; Nelson et al., 2020), loss-of-function mutations in *hip14* cause increased acoustic startle sensitivity and defects in acoustic startle habituation (**Fig.1C-D**).

**Figure 1.**
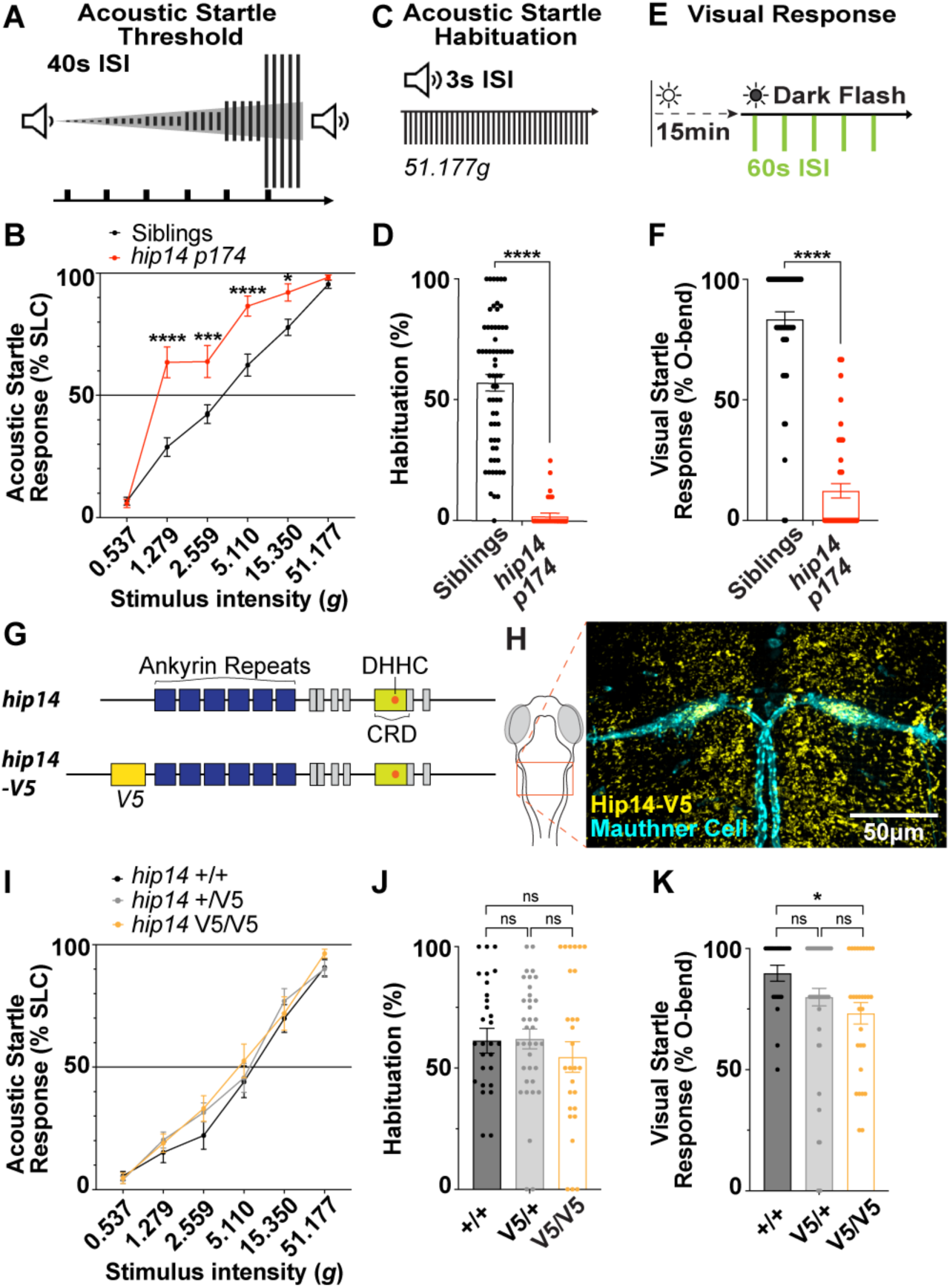
V5-knockin enables detection of endogenous, functional Hip14 expression and dynamics. (**A**) Schematic of assay for acoustic startle thresholds (**B**) *hip14^p174^*mutants are hypersensitive to acoustic stimuli. Differences between siblings (n=69) and mutants (n=38) are significant at stimulus intensities 1.279*g*, 2.559*g*, 5.11*g*, and 15.350*g* (\*\*\*\**p* <0.0001, \*\*\**p*=0.0008, \*\*\*\**p* <0.0001, \**p*=0.0452) via Šídák Two-way ANOVA. **(C)** Schematic of assay for acoustic startle habituation. **(D)** Habituation is significantly disrupted in *hip14* mutants, n=34, relative to siblings, n=64 (Mann-Whitney test, \*\*\*\**p*<0.0001). (**E**) Assay schematic for dark flash. (**F**) *hip14^p174^* mutants (n=59) display reduced responses to visual stimuli relative to their siblings (n=49), Mann-Whitney test (\*\*\*\**p*<0.0001). (**G**) Schematic indicating the site of the *hip14-V5* insertion and (**H**) endogenous Hip14-V5 expression in the hindbrain (yellow=Hip14-V5, cyan=*UAS:Gap43:Citrine* expressed under the control of the *GFFDMC130a gal4* in the Mauthner cell). (**I-K**) V5 insertion into the *hip14* locus does not alter acoustic startle behaviors and only subtly impacts visual responses. (**I**) Differences between wild-type (*hip14*^+/+^) n=31, heterozygotes (*hip14^+/V5^*) n=44, and mutants (*hip14^V5/V5^*) n=32 are not significant, Two-way ANOVA with Tukey’s test for multiple comparisons. From the lowest to highest stimulus intensities, the differences between wild-type (*hip14*^+/+^) and mutants (*hip14^V5/V5^*) are not significant (*p*=0.9909, 0.8553, 0.2486, 0.4245, 0.9514, and 0.6764, respectively). (**J**) Acoustic startle habituation was analyzed with one-way ANOVA with Tukey’s test for multiple comparisons and yielded no significant differences when comparing wild type (*hip14*^+/+^), n=29, with heterozygotes (*hip14^+/V5^*), n=36, *p*=0.9942, and mutants (*hip14^V5/V5^*), n=29, *p*=0.6429. (**K**) Dark flash assay analyzed with non-parametric Kruskal-Wallis test on wild-type (*hip14*^+/+^) n=21, heterozygotes (*hip14^+/V5^*) n=56, and mutants (*hip14^V5/V5^*) n=30, and \**p*=0.0276 for *hip14*^+/+^ comparison with *hip14^V5/V5^*.

In addition to the acoustically evoked startle, zebrafish perform large angle turns when subjected to sudden darkness, or dark flash (DF) stimuli. The DF response relies on distinct circuitry relative to the acoustic startle (Burgess and Granato, 2007b; Randlett et al., 2019). To determine whether Hip14 function also contributes to visual startle behaviors, we performed a DF assay, in which we light-adapted 6 dpf larvae and then subjected them to 5 DF stimuli separated by a 60s ISI (**Fig.1E**) (Burgess and Granato, 2007b; Wolman et al., 2011, 2014; Randlett et al., 2019). We found that *hip14^p174^*mutants showed a significantly reduced DF response (\*\*\*\**p*<0.0001) compared to their siblings (**Fig.1F**), indicating that in addition to regulating startle sensitivity and promoting acoustic startle habituation, Hip14 functions to promote detection of or responsiveness to visual stimuli.

### Hip14-V5 reveals endogenous expression in multiple cell types of the acoustic startle circuit

Prior work suggests that PAT subcellular localization is a key determinant of substrate palmitoylation (Globa and Bamji, 2017). In *Drosophila*, Hip14 localizes to presynapses and becomes re-localized to the plasma membrane when synaptic vesicle endocytosis is blocked (Ohyama et al., 2007; Stowers and Isacoff, 2007). These data prompted us to hypothesize that Hip14 could localize to distinct subcellular compartments within cells of the acoustic startle circuit, and that its subcellular localization could inform its function. To assess this, we first set out to develop a tool to examine endogenous expression of Hip14 within cells of the acoustic startle circuit that would additionally allow us to assess potential dynamics in its sub-localization. For this, we used CRISPR-Cas9 genome editing to insert a V5 epitope tag in-frame into the *hip14* coding sequence and visualized endogenous Hip14 protein via IHC using antibodies against V5 (**Fig.1G-H**). Consistent with, and expanding on prior work on *hip14* gene expression that showed high enrichment in the central nervous system (CNS) in adult mice (Singaraja et al., 2002; Huang et al., 2004; Wild et al., 2022) and zebrafish (Shi et al., 2015), we found Hip14-V5 protein expression throughout the brain and spinal cord of larval zebrafish. In particular, we document Hip14-V5 protein in the M-cells (**Fig.1H**), the reticulospinal neurons responsible for driving the initial phase of the short-latency acoustic startle response in larval zebrafish (Kimmel et al., 1980; Korn and Faber, 2005; Burgess and Granato, 2007a). Next, to ensure that the *V5* insertion does not disrupt the function of Hip14, we intercrossed heterozygous carriers of the *V5* insertion to obtain larvae in which both copies of *hip14* contain the in-frame V5 tag (*hip14^V5/V5^)*. Subsequently, we conducted the acoustic startle sensitivity, habituation, and dark flash assays on homozygous knock-in larvae and compared their response profiles to their siblings carrying one or zero copies of the *V5* knock-in (*hip14^+/V5^ and hip14^+/+^,* respectively). We found that *hip14^V5/V5^* animals do not exhibit any significant differences in acoustic startle sensitivity or habituation (**Fig.1I-J**). Dark flash responses are slightly affected when comparing wild-type *hip14^+/+^* to *hip14^V5/V5^* (\**p*=0.0276, **Fig.1K**). Overall, our results confirmed that *hip14*-*V5* epitope tag knock-in does not affect Hip14 function within the acoustic startle circuit. Altogether, our new *hip14-V5* knock-in enables the detection of endogenous functional Hip14 expression, localization, and dynamics.

Next, using this new tool, we examined endogenous expression of Hip14 within individual components of the acoustic startle circuit. We found that in addition to its expression throughout the nervous system, including in M-cells (**Fig 2A-D**), Hip14 is expressed in mechanosensory hair cells in the inner ear that detect acoustic/vibrational stimuli (**Fig 2E-H**).

**Figure 2.**
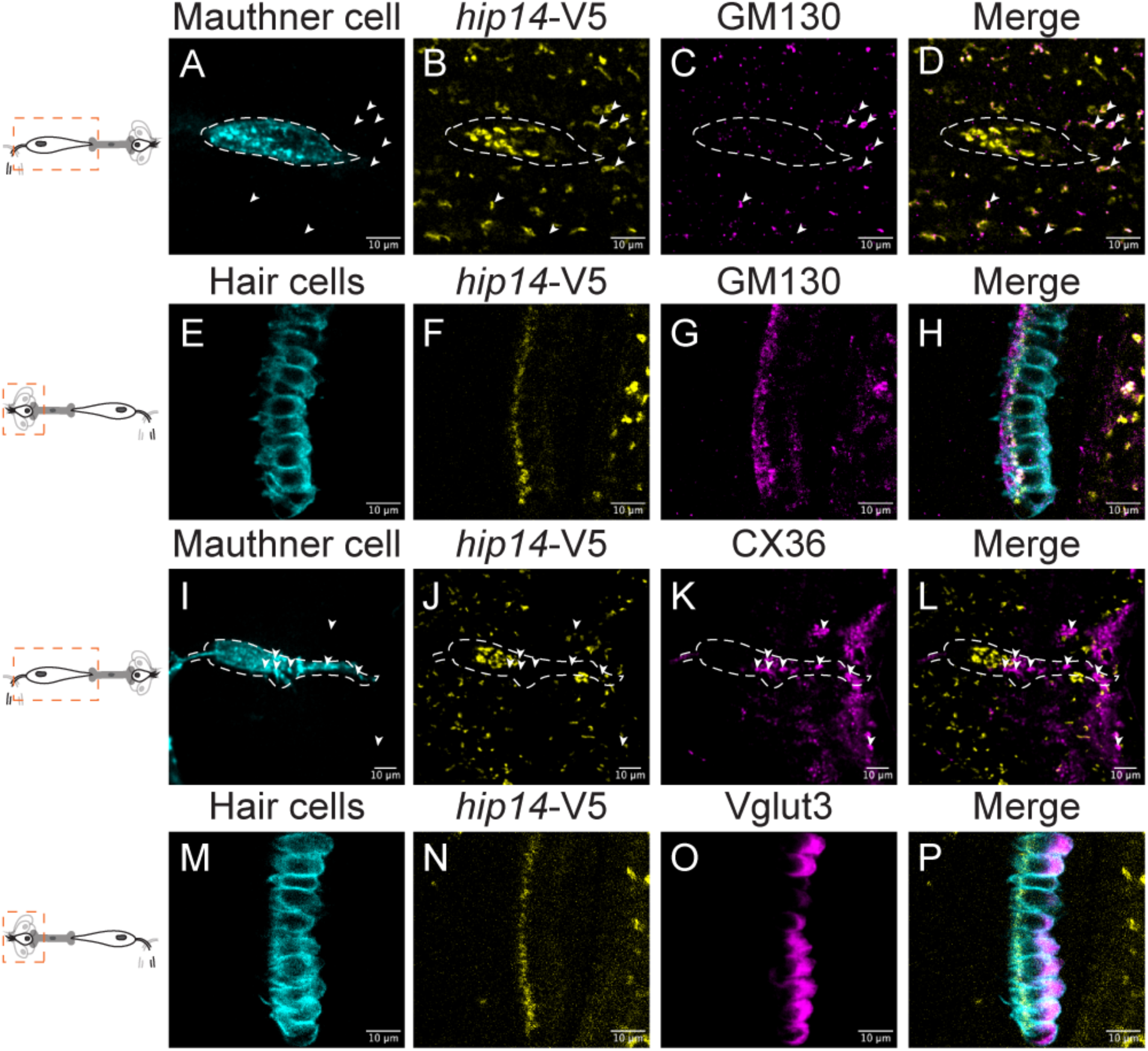
Hip14 is localized to the Golgi apparatus in the M-cell and in mechanosensory hair cells but does not localize to presynaptic terminals. (**A-D**) Single plane confocal images of M-cells labeled with **(A)** *GFFDMC130a; UAS:Gap43:Citrine* marker (cyan), **(B)** Hip14-V5 (yellow) **(C)** tdTomato-GM130 (magenta), **(C)** Merge; arrowheads indicate examples of co-localization of tdTomato-GM130 and Hip14-V5 outside the M-cell, which is encircled by a dashed line **(E-H)** Mechanosensory hair cells of the saccular macula labeled with **(E)** *myo6b:GCaMP6s:CAAX* (cyan). **(F-H)** same as **B-D; (I, J, L)** Same as A, B. **(K)** Gap junctions labeled with Cx36. **(L)** Merge. Arrowheads indicate Cx36 puncta, which do not colocalize with Hip14-V5. Dashed line encircles M-cell. **(M, N)** Same as **E, F. (O)** Saccular hair cell presynaptic vesicle pools labeled with Vglut3 antibody **(P)** Merge.

### Hip14 localizes to the Golgi apparatus within mechanosensory hair cells and neurons of the acoustic startle circuit

Subcellularly, Hip14 is best known to localize to the cis-Golgi where it palmitoylates a variety of substrates (Singaraja et al., 2002; Huang et al., 2004; Ohno et al., 2006; Ernst et al., 2018; Nthiga et al., 2021). However, the *Drosophila* HIP14 ortholog (dHIP14) was also found to be enriched at presynaptic sites, where it co-localizes with CSP and synaptobrevin (Ohyama et al., 2007; Stowers and Isacoff, 2007). Furthermore, when synaptic vesicle endocytosis from the plasma membrane was disrupted, dHIP14 re-localized to the plasma membrane, indicating that it is potentially present on synaptic vesicles (Ohyama et al., 2007). Since habituation is thought to involve short-term plasticity, particularly at presynaptic sites (Weber et al., 2002; Simons-Weidenmaier et al., 2006; Marsden and Granato, 2015), we hypothesized that Hip14 would exhibit a presynaptic localization within neurons of the acoustic startle circuit in larval zebrafish. To test this hypothesis, we used the *hip14^V5^*reporter together with a transgenic line that we generated to label the *cis* Golgi apparatus in all cells (*p24b.ubi:tdTomato-GM130*) (**Fig. 2A-H**). We found that throughout the hindbrain, Hip14-V5 puncta co-localized with the tdTomato-GM130 marker. While in the M-cell itself, the *ubi:tdtomato-GM130* transgene failed to reach detectable levels of expression, the Hip14-V5 puncta were localized to the perinuclear region (**Fig.2A-D**), consistent with Hip14 localization to the Golgi apparatus within the M-cell (**Fig. 2A-D**). Next, we examined Hip14-V5 subcellular localization in mechanosensory hair cells of the inner ear. Specifically, we focused on the saccule, an otolith organ considered the primary site of sound detection in larval zebrafish (Lu and DeSmidt, 2013; Yao et al., 2016; Favre-Bulle et al., 2020). Mechanosensory hair cells are polarized, with pre-synaptic specializations forming basally and stereocilia extending from the apical pole. We found that Hip14 subcellular localization is also polarized: Hip14 is present at the apical pole but not at the basal pole (**Fig.2E-H**). Similarly, the tdTomato-GM130 reporter localized apically and was largely absent from the basal pole (**Fig. 2E-H**). Together, our data indicate that Hip14 predominantly localizes to the Golgi apparatus, akin to many other PATs (Globa and Bamji, 2017; Philippe and Jenkins, 2019), in neurons and hair cells comprising the acoustic startle circuit.

M-cells receive mixed chemical and electrical synapses from the VIIIth nerve at the level of the lateral dendrite (Pereda et al., 2004; Yao et al., 2014). To examine whether Hip14 localizes to the presynaptic terminals of VIIIth nerve afferents onto the M-cell, we used antibodies against Connexin 36 (Cx36), which recognizes zebrafish Connexin homologs present at VIIIth nerve – M-cell synapses (including Cx34.7, Cx35.1, Cx35.5, and Cx34.1) (Rash et al., 2013; Miller et al., 2017). We again found that Hip14 does not co-localize with CX36 gap-junction aggregates apposed to the lateral dendrite (**Fig. 2I-L**), leading us to reject the hypothesis that Hip14 localizes to presynaptic sites targeting the M-cell lateral dendrite. Next, to rule out presynaptic localization of Hip14 in mechanosensory hair cells, we stained for the Vesicular Glutamate Transporter 3 (Vglut3), which localizes to synaptic vesicle pools organized at the basal pole of each mechanosensory hair cell (Obholzer et al., 2008). We found that these basally located synaptic vesicle pools do not overlap with the apically localized Hip14-V5 (**Fig. 2M-P**). To further determine whether Hip14 is presynaptically localized in other regions of the CNS, we visualized Hip14 expression relative to synaptic neuropil throughout the brain by generating a transgenic line *NeuroD:SypB-mScarlet* that labels the presynaptic vesicle protein Synaptophysin b (SypB) (Hughes and Appel, 2019). When we examined Hip14 expression in animals carrying this presynaptic reporter, we found that, as before, Hip14 is broadly expressed throughout the brain. Yet, in the optic tectum neuropil, a region densely packed with synaptic connections (Stuermer, 1988) and strongly labeled by SypB-mScarlett, we observed a relative void in the Hip14-V5 signal, with only rare Hip14-V5 puncta. Notably, the few Hip14-V5 puncta in this region were present in voids lacking the SypB-mScarlett reporter. (**Supplemental Fig.S1A**). Overall, our results demonstrate that Hip14 localizes predominantly to the Golgi apparatus and is absent from presynaptic regions, leading us to a model in which Hip14 functions at the level of the Golgi and not directly at synapses to control sensory-evoked behaviors.

### Pan-neuronal expression of Hip14 rescues acoustic startle threshold, habituation, and dark-flash responses in *hip14* mutants

Given that Hip14 is endogenously expressed both in peripheral sensory cells and neurons of the acoustic startle circuit, we next tested whether Hip14 functions within neurons or within mechanosensory hair cells to regulate behavior. For this, we inserted into the *pIGLET14a* safe-harbor locus (Lalonde et al., 2024), a transgene expressing Hip14-EGFP under the control of the pan-neuronal *HuC* promoter (Kim et al., 1996; Park et al., 2000; Higashijima et al., 2003): *p14a.Huc:hip14-egfp* (**Fig.3A)**. Pan-neuronal overexpression of Hip14-EGFP fully rescued acoustic startle habituation deficits in *hip14^p174^*mutants (\*\*\*\**p*<0.0001) (**Fig. 3B**). Pan-neuronal overexpression of Hip14-EGFP in *hip14^p174^* mutants similarly restored startle responses to visual DF stimuli (**Fig. 3C**) (\*\*\*\**p*<0.0001). For acoustic startle behaviors, pan-neuronal overexpression of Hip14 fully rescued the hypersensitivity of *hip14* mutant animals to sibling levels, while having no effect on siblings (**Fig.3D**) (At stimulus intensities of 1.279*g*, 2.559*g*, 5.11*g*, and 15.35*g*, the Hip14 transgene significantly, \*\*\*\**p*<0.0001, reduced the probability of response in *hip14^p174^* mutants, Odds Ratio (OR): 0.1164, 0.0639, 0.0979, and 0.0623, respectively). Together, these data are consistent with a model in which Hip14 functions in neurons to regulate acoustic and visual startle behaviors.

**Figure 3.**
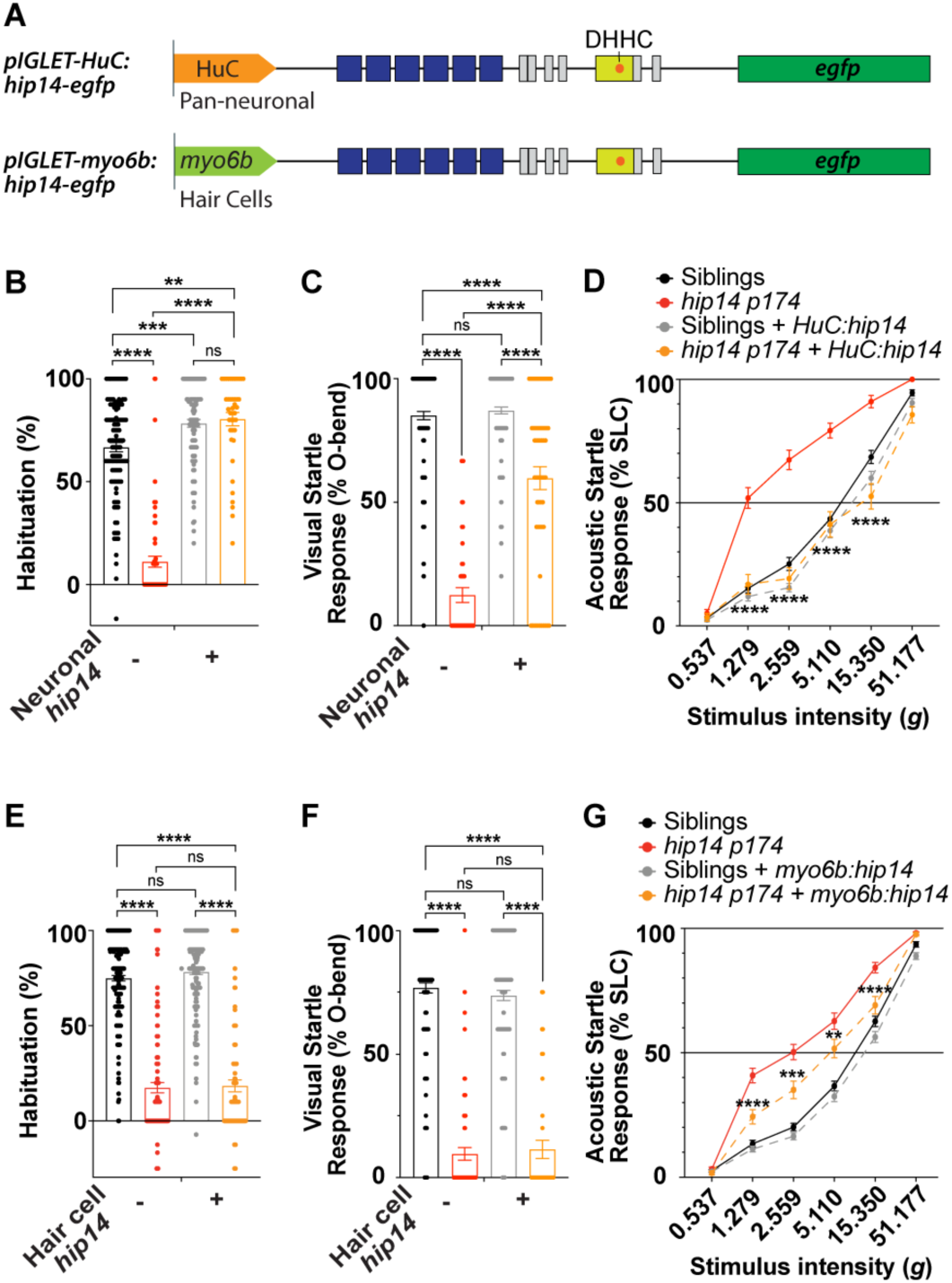
Pan-neuronal expression of Hip14 restores acoustic and visual behaviors, while hair cell expression of Hip14 partially restores acoustic startle sensitivity. *p14a.Huc:hip14-egfp* and *p14a.myo6b:hip14-egfp* were expressed in the *hip14 ^p174^* mutant background. All statistical analyses here used Two-way ANOVA with Tukey’s multiple comparison tests, except acoustic startle sensitivity assays **(D, G)**, which were analyzed using Logistic Mixed-Model Estimated Marginal Means pairwise tests (LMM) with the Benjamini-Hochberg procedure. **(A)** Schematic of pan-neuronal (*HuC*) and mechanosensory hair cell (*myo6b*) driven *hip14*-*egfp* transgenes. **(B)** Mutants (n=68) displayed significantly lower acoustic startle habituation than their siblings (n=112) (predicted mean difference: 55.69, \*\*\*\**p*<0.0001), whereas pan-neuronal expression of *hip14-egfp* in mutants (n=41) rescued habituation (predicted mean difference: 69.36, \*\*\*\**p*<0.0001). Siblings with transgene (n=105) and mutants with transgene (n=41) exhibited a slight increase compared with siblings (\*\*\**p*=0.0006, \*\**p*=0.0036, respectively). **(C)** Pan-neuronal expression of *hip14-egfp* partially rescued dark flash response (predicted mean difference: 72.69, \*\*\*\**p*<0.0001), siblings (n=158), mutants (n=47), siblings+*hip14-egfp* (n=163), mutants + *hip14-egfp* (n=58). **(D)** Pan-neuronal expression of *hip14-egfp* rescued acoustic startle hypersensitivity of mutants (LMM: \*\*\*\**p*<0.0001 at stimulus intensities of 1.279*g*, 2.559*g*, 5.11*g*, and 15.350*g*, respectively). Mutants + *hip14-egfp* (n=46) and mutants without transgene (n=71) were compared to siblings (n=123) and siblings + *hip14-egfp* (n=118). **(E)** Hair cell overexpression of *hip14-egfp* in mutants (n=105) failed to rescue the habituation defects of mutants (n=141) (*p*=0.9894). Siblings (n=217) and siblings with the transgene (n=215) showed no significant difference (*p*=0.5309). **(F)** Hair cell overexpression of *hip14-egfp* in mutants (n=40) failed to rescue the initial DF response of mutants (n=65) (*p*=0.9855). Siblings (n=170) and siblings + *myo6b:hip14-egfp* (n=171) showed no significant difference (*p*=0.6942). **(G)** Hair cell overexpression of the *hip14-egfp* transgene in mutants (n=152) partially rescued hypersensitivity of mutants (LMM: **** *p*<0.0001, \*\*\**p*=0.000157, \*\**p=* 0.00412, at stimulus intensities of 1.279*g*, 2.559*g*, 5.11*g*, and 15.350*g*). Mutants with transgene (n=152) and mutants without transgene (n=111) were compared to siblings (n=267) and siblings with transgene (n=251).

Next, since we found that Hip14 is also expressed in inner ear hair cells (**Fig.2E-H**), we asked whether it modulates acoustically evoked behaviors at the level of sound detection within these peripheral sensory cells. For this, we generated a transgenic line in the same safe harbor locus (*p14a.myo6b:hip14-egfp*) that drives expression of Hip14 in all mechanosensory hair cells (Obholzer et al., 2008). As expected, *hip14-egfp* expression in hair cells did not restore habituation or responses to visual stimuli (**Fig.3E-F**). These data are consistent with our understanding of habituation as a short-term learning process that relies on neuronal mechanisms (Rankin et al., 2009), and dark flash responses as reliant on sensory detection within the retina but not the inner ear. Interestingly, we found that *hip14-egfp* expression in hair cells partially rescued acoustic hypersensitivity in the *hip14^p174^*mutant (Reduced response odds in mutants at stimulus intensity of 5.11*g,* OR=0.482, \*\**p*=0.00412) compared to mutants without the *hip14-egfp* transgene (**Fig.3G**). Therefore, both pan-neuronal and hair cell-specific overexpression of Hip14 rescued acoustic startle thresholds, although to different extents (**Fig.3B,E**). Our data are consistent with a model in which excitable cells, including both mechanosensory hair cells and neurons, are broadly hyperactive in *hip14* mutants. Reintroduction of Hip14 within either cell type is enough to limit hyperexcitability within the circuit, damping the acoustic hypersensitivity phenotype. Together, our results establish that Hip14, although it functions primarily in neurons, also acts partially in hair cells to regulate acoustic startle thresholds.

### Hip14 controls behavior in part through a mechanism independent of its palmitoyl acyltransferase (DHHC) domain

To better understand how Hip14 controls behavior, we set out to test the functional requirements of each of its protein domains. First, to understand the role of the DHHC domain, we generated a *hip14* transgene with a mutated DHHC domain, substituting the catalytic cysteine with serine, thereby abolishing its auto-acylation and rendering the domain catalytically inactive (Lobo et al., 2002) (**Fig.4A**). We then integrated the transgene into the same *p14a* safe-harbor locus that was used for our WT pan-neuronal (*p14a.Huc:hip14-egfp)* and hair cell *hip14* rescue constructs (*p14a.myo6b:hip14-egfp*) to enable precise comparison of rescue efficiency across experiments. Then we conducted acoustic startle and visual sensitivity as well as acoustic startle habituation experiments as before. The catalytically inactive Hip14-DHHS transgene did not restore normal habituation. Surprisingly, neuronal expression of the Hip14-DHHS transgene was sufficient to fully rescue acoustic startle response thresholds (**Fig.4A,B**) (Transgene reduced the odds of mutants responding at stimulus intensities 1.279*g* (\*\**p*=0.0014, OR 0.1688), 2.2559*g* (\*\*\*\**p*<0.0001, OR 0.0596), 5.11*g* (\*\*\**p*= 0.0003, OR 0.1211), and 15.35*g* (\**p*= 0.0268, OR 0.1810)). These data are consistent with a model in which the Hip14 DHHC domain is required for acoustic startle habituation but dispensable for regulating innate thresholds and provides a clear genetic separability for the often-linked phenotypes of behavioral hypersensitivity and inability to habituate.

**Figure 4.**
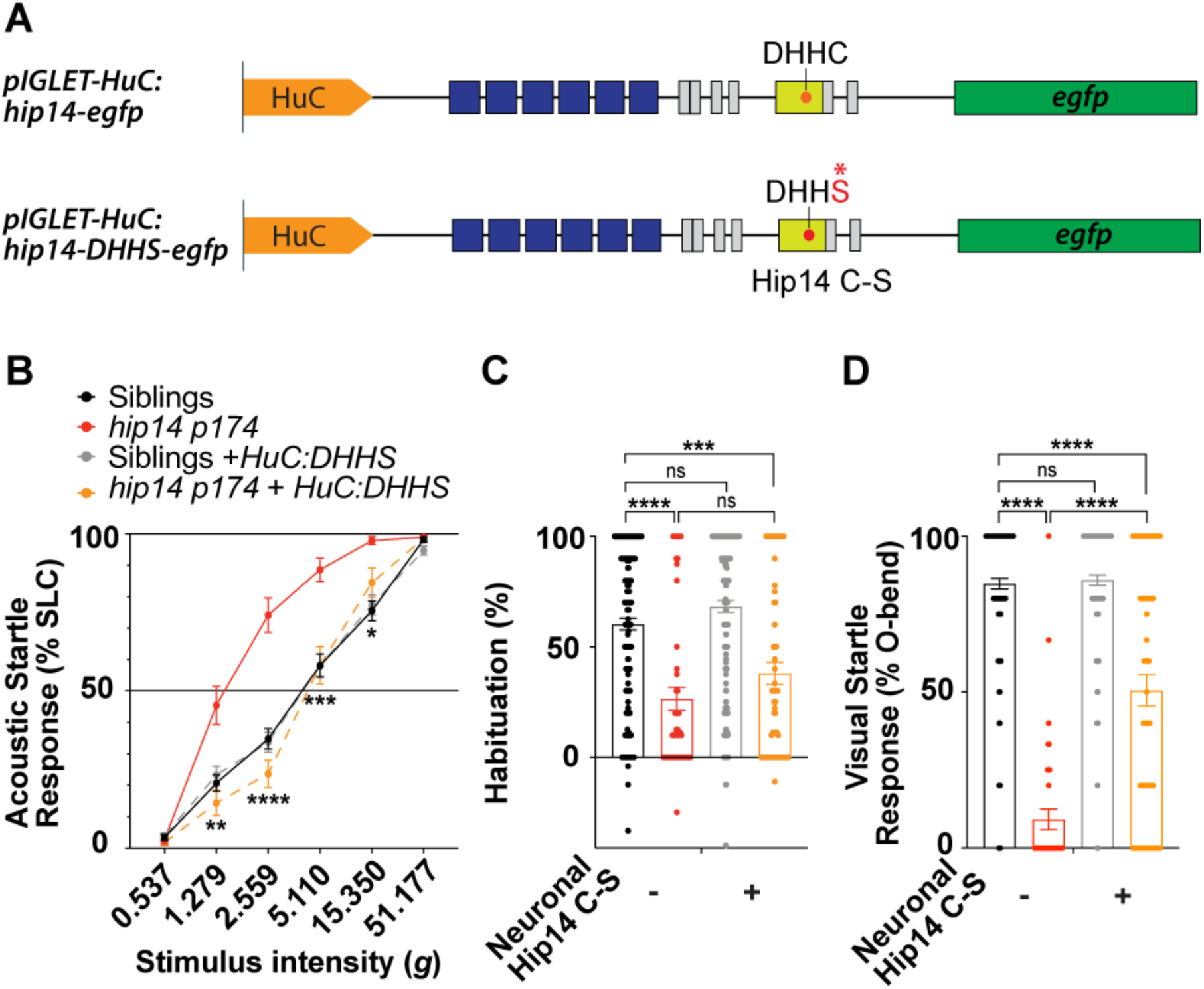
Catalytically inactive Hip14 transgene (DHHS) rescues acoustic startle sensitivity and dark flash responses but failed to rescue acoustic startle habituation. We expressed the catalytically inactive *p14a.Huc:hip14-DHHS-egfp* transgene in the *hip14 ^p174^* mutant background. All statistical analyses, unless specified otherwise, used Tukey’s multiple comparison tests. **(A)** Schematics of pan-neuronal (*HuC*) driven WT and *hip14*-*DHHS* (C-S) transgenes. (**B)** Pan-neuronal expression of *hip14-DHHS-egfp* fully rescued hypersensitivity of mutants (Logistic Mixed-Model Estimated Marginal Means pairwise tests (LMM) with the Benjamini-Hochberg procedure: \*\**p*=0.00137, \*\*\*\**p*<0.0001 at stimulus intensities of 1.279*g*, 2.559*g*, 5.11*g*, and 15.350*g*). Mutants with transgene (n=48) and mutants without transgene (n=43) were compared to siblings (n=129) and siblings with transgene (n=137). (C) *hip14-DHHS* transgene failed to rescue acoustic habituation of *hip14* mutants (*p*=0.3279); siblings n=184, mutants n=53, siblings+*hip14-DHHS* n=169, mutants + *hip14-DHHS* n=64. (**D**) Pan-neuronal *hip14-DHHS-egfp* in mutants (n=66) partially rescued the dark flash response of *hip14* mutants (n=40). Predicted mean difference of 41.27%,\*\*\*\**p*<0.0001. Siblings (n=188) and siblings with the transgene (n=170) show no differences in percent response, *p*=0.9750.

Next, we examined visual responses. While *hip14* mutants showed a lower probability of performing DF responses as expected, (**Fig. 4D**) overexpressing the DHHS transgene in mutants significantly rescued the response rate of *hip14* mutants, from 9.208% to 50.48% (Two-way ANOVA Tukey, \*\*\*\**p*<0.0001), but not to the level observed in siblings (84.76%, \*\*\*\**p*<0.0001, **Fig. 3D).** Nonetheless, pan-neuronal expression of the *hip14-DHHS-egfp* rescues visual response rates in mutants (predicted mean difference 41.27%, \*\*\*\**p*<0.0001 **Fig.4D**) to a similar extent as compared to overexpression of the WT *hip14* transgene (predicted mean difference 47.33%, \*\*\*\**p*<0.0001 **Fig.3D**). Taken together, we find that the DHHC domain is dispensable for promoting responses to visual stimuli and limiting sensitivity to acoustic stimuli. These data also suggest that additional molecular functions of Hip14, aside from DHHC-dependent palmitoylation, are relevant *in vivo*.

### Hip14 ankyrin repeat domains are required for both establishment of sensory thresholds and habituation

Aside from the palmitoylation (DHHC) domain, Hip14 also possesses six Ankyrin repeats, which facilitate protein-protein interactions, including with its own palmitoylation substrates (Lemonidis et al., 2014). To assess the contribution of the AR domains to Hip14 regulation of behavior, we generated a Hip14 rescue construct in which the ankyrin repeats are deleted (*p14a.HuC:hip14-ΔAR-egfp*, **Fig.5A**), which has previously been shown to abolish many substrate interactions (Lemonidis et al., 2014; Butler et al., 2023). We overexpressed *hip14-ΔAR-egfp* under the control of the pan-neuronal *HuC* promoter in *hip14^p174^* mutants. *hip14-ΔAR-egfp* failed to rescue acoustic startle habituation and DF basal response. On the other hand, pan-neuronal overexpression of this construct induced a significant but partial rescue of acoustic hypersensitivity relative to *hip14^p174^*mutants without the transgene (**Fig.5B-D)**. At stimulus intensities of 1.279*g*, 2.2559*g*, and 5.11*g,* mutants with the ΔAR transgene all showed significant (\*\*\*\**p*<0.0001) reductions in response odds, yielding odds ratios of 0.403, 0.402, and 0.379, respectively. These data document that even without the AR domain, Hip14 still retains partial ability to regulate acoustic startle thresholds.

**Figure 5.**
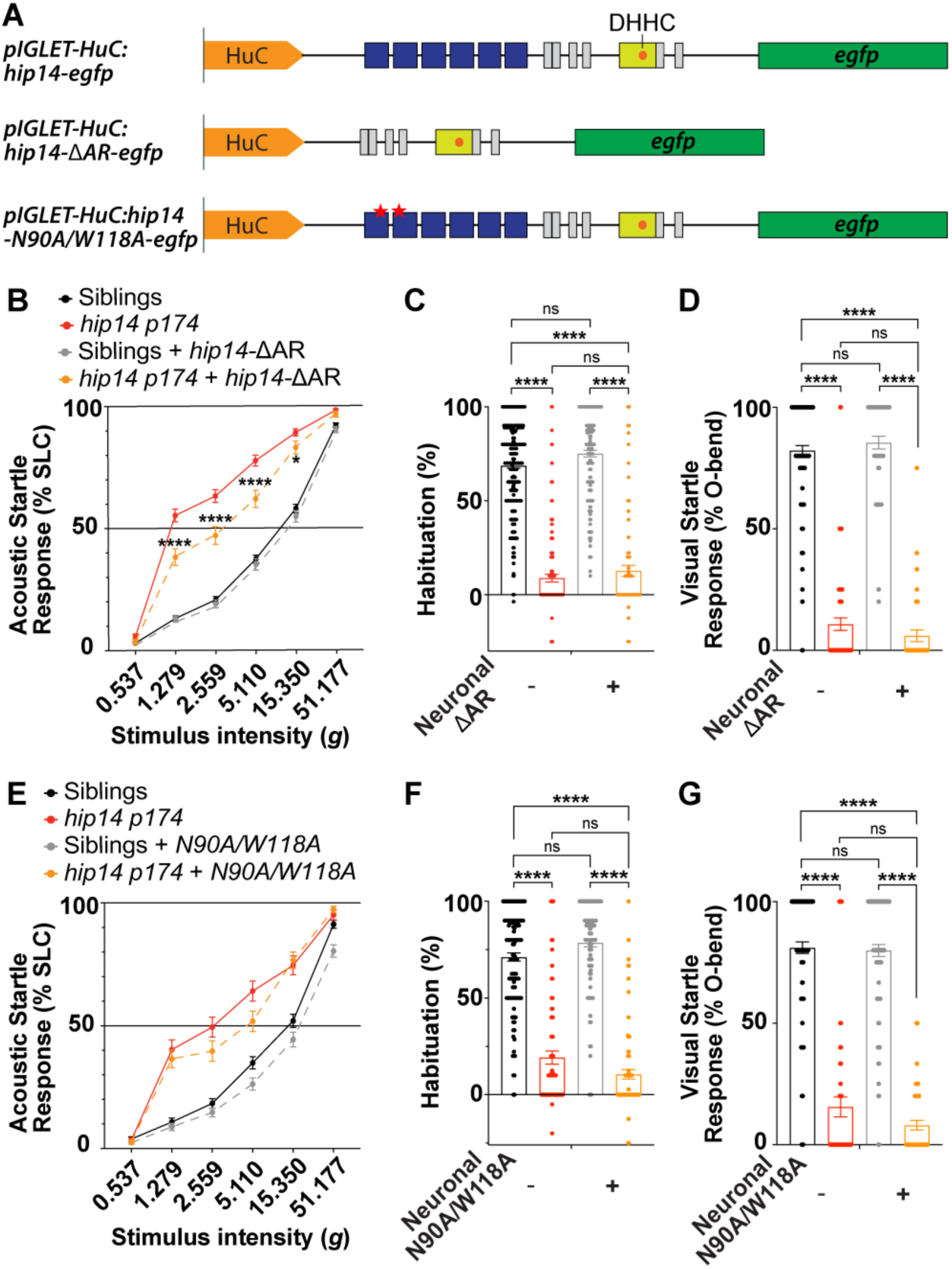
Transgenes lacking the Hip14 ankyrin repeats fail to rescue acoustic startle habituation or dark flash, and only partially rescue acoustic startle sensitivity. **(A)** Schematics of pan-neuronal (*HuC*) driven *hip14* AR deletion (ΔAR) versus point mutation (N90A/W118A) transgenes. All statistical analyses here used Two-way ANOVA with Tukey’s multiple comparison tests, except acoustic startle sensitivity assays **(B, E)**, which were analyzed using Logistic Mixed-Model Estimated Marginal Means pairwise tests (LMM) with the Benjamini-Hochberg procedure. **(B-D)** Pan-neuronal expression of *hip14-ΔAR-egfp*. **(B)** Pan-neuronal expression of *hip14-ΔAR* partially rescues acoustic startle thresholds (LMM: \*\*\*\**p*<0.0001, \**p*=0.0151 at stimulus intensities of 1.279*g*, 2.559*g*, 5.11*g*, and 15.350*g*), siblings (n=368), mutants (n=195), siblings+*hip14-ΔAR* (n=188), mutants+*hip14-ΔAR* (n=109). **(C)** Pan-neuronal *Hip14-ΔAR* failed to rescue acoustic startle habituation in mutants (*p*=0.6103; siblings n=331, mutants (n=187), siblings+*hip14-ΔAR* (n=174), mutants +*hip14-ΔAR* (n=102)). **(D)** *Hip14-ΔAR* transgene failed to rescue DF response (*p*=0.6479; siblings (n=115), mutants (n=70), siblings+*hip14-ΔAR* (n=54), mutants+*hip14-ΔAR* (n=39). **(E-G)** Pan-neuronal expression of *hip14-AR* point-mutation transgene (*N90A/W118A*) did not alter acoustic or visual behaviors. **(E)** N90A/W118A transgene did not rescue hypersensitivity in mutants (LMM: *p*=0.5990, 0.5990, 0.1264, 0.06343, 0.6829, and 0.3374, from the lowest to highest stimulus intensities); siblings (n=145), mutants (n=81), siblings+*N90AW118A* (n=121), mutants+*N90A/W118A* (n=85). **(F)** Pan-neuronal *N90A/W118A* failed to rescue acoustic startle habituation in mutants (*p*=0.1234; siblings n=136, mutants n=75, siblings+*N90AW118A* n=109, mutants+N90A/W118A n=81). **(G)** Pan-neuronal *N90A/W118A* failed to rescue DF response in mutants (*p*=0.4143; siblings n=116, mutants n=55, siblings+*N90A/W118A* n=95, mutants+*N90A/W118A* n=47).

Human ZDHHC17 was previously identified to have two critical residues in the ankyrin repeat domains: Asparagine 100 and Tryptophan 130; both are crucial for the interaction between Hip14 and its palmitoylation substrate, SNAP25 (Verardi et al., 2017; Butler et al., 2023). Alanine mutagenesis of critical residues, N100 and W130, was found to abolish the interaction *in vitro* (Verardi et al., 2017). Similarly, N100A/W130A mutations in HeLa cells abolished the zDHHC17 interaction with Calcoco1a and Tax1bp1 (Nthiga et al., 2021). To further distinguish whether the partial rescue by ΔAR was specifically caused by a disrupted zDABM-ligand interface that has been previously identified (Verardi et al., 2017; Nthiga et al., 2021; Butler et al., 2023), we generated Alanine mutations in the residues corresponding to A100 and W130 (zebrafish sequence: N90A and W118A) within the ankyrin repeat region: *p14a.HuC:hip14-N90A/W118A-egfp* (**Fig.5A)**. We found that transgenes carrying these point mutations in the AR, although trending toward rescue at stimulus intensities of 1.279g, 2.559g, and 5.11g, did not produce statistically significant rescue (*p*=0.599, 0.1264, and 0.0634, respectively) (**Fig.5E**). Furthermore, neither habituation nor DF response displayed any trend toward rescue as in the case of the full AR deletion transgene (**Fig.5F,G**). Together, our data support a model in which ankyrin repeats are essential for Hip14 function and indicate that only limited functionality is retained when the ankyrin repeats are removed.

## Discussion

To regulate attention to environmental cues, animals tune their sensitivity to stimuli (Burgess and Granato, 2007a; Wolman et al., 2011; Marsden et al., 2018; Ogino et al., 2019). Innate thresholds are established during development, and habituation enables animals to tune thresholds based on previous sensory experiences (Simons-Weidenmaier et al., 2006; Burgess and Granato, 2007a; Rankin et al., 2009; Roberts et al., 2011; Wolman et al., 2011; Lau et al., 2013; Barron et al., 2017, 2017). The zebrafish acoustic startle response provides an ideal model to understand the molecular and cellular mechanisms through which innate acoustic startle thresholds are established and undergo plasticity. We previously found that the palmitoyl acyltransferase, Hip14, functions both to promote acoustic startle response thresholds and to promote their plasticity through habituation (Nelson et al., 2020, 2023). Here, we explored the cellular and molecular mechanisms through which Hip14 functions.

### Cellular mechanisms underlying the establishment of innate acoustic startle response thresholds are distinct from those underlying acoustic startle response habituation

The mechanisms underlying establishment of acoustic startle response thresholds are thought to be closely linked to those regulating habituation. Genetic mutations and pharmacological perturbations that induce habituation deficits are often also associated with lowered innate startle response thresholds (Wolman et al., 2011, 2015; McDiarmid et al., 2020; Nelson et al., 2020; Santistevan et al., 2021; Zúñiga Mouret et al., 2024; Schloss et al., 2025). It is plausible, therefore, that in some cases, defects in acoustic habituation partially or fully result from reduced acoustic startle thresholds. Supporting this view, hypersensitive *hip14* mutants can habituate if stimulus intensity is lowered, although at a reduced rate relative to their WT siblings (Nelson et al., 2023). On the other hand, acoustic startle hypersensitivity is occasionally observed in the absence of habituation deficits (Marsden et al., 2018; Ortiz et al., 2024).

Our results demonstrate that even when acoustic startle habituation and acoustic startle hypersensitivity co-occur, they rely on distinct molecular mechanisms. We found that a catalytically inactive version of the *hip14* transgene (*hip14-DHHS*) rescued acoustic startle thresholds but failed to rescue acoustic startle habituation. These data genetically distinguish innate startle threshold establishment from habituation: in animals carrying the transgene, thresholds are normal, but habituation deficits remain. They further indicate that Hip14-dependent palmitoylation is critical for habituation learning, whereas sensory threshold establishment relies on a distinct mechanism (**Fig.3**).

Our results not only support the idea that sensory thresholds and habituation rely on distinct cellular processes but also show that the palmitoylation via the DHHC domain, although often seen as Hip14’s primary function, is not required for all facets of its molecular function.

### Key residues in the ankyrin repeat domains are essential for nearly all Hip14 functions

The Hip14 AR domain enables interactions with many downstream interactors, including SNAP25, CSP (Lemonidis et al., 2014, 2015), Htt (Huang et al., 2011), and JNK (Yang and Cynader, 2011). In most circumstances, Hip14 AR-dependent interactions support substrate palmitoylation (Lemonidis et al., 2014, 2015; Verardi et al., 2017), though exceptions exist in which Hip14 still induces substrate palmitoylation without these key domains (Hemsley and Grierson, 2011). Here, we found that the *hip14*-ΔAR transgene partially rescued acoustic startle thresholds but failed to rescue acoustic startle habituation or visual responsiveness (**Fig. 5B**). Together with our finding that *hip14-DHHS* rescues acoustic startle thresholds (**Fig. 4B**), we propose that the Ankyrin repeat domains are largely required for the role of Hip14 in promoting normal behavior, while the DHHS domain is required specifically for habituation (**Fig.6A,B**).

**Figure 6.**
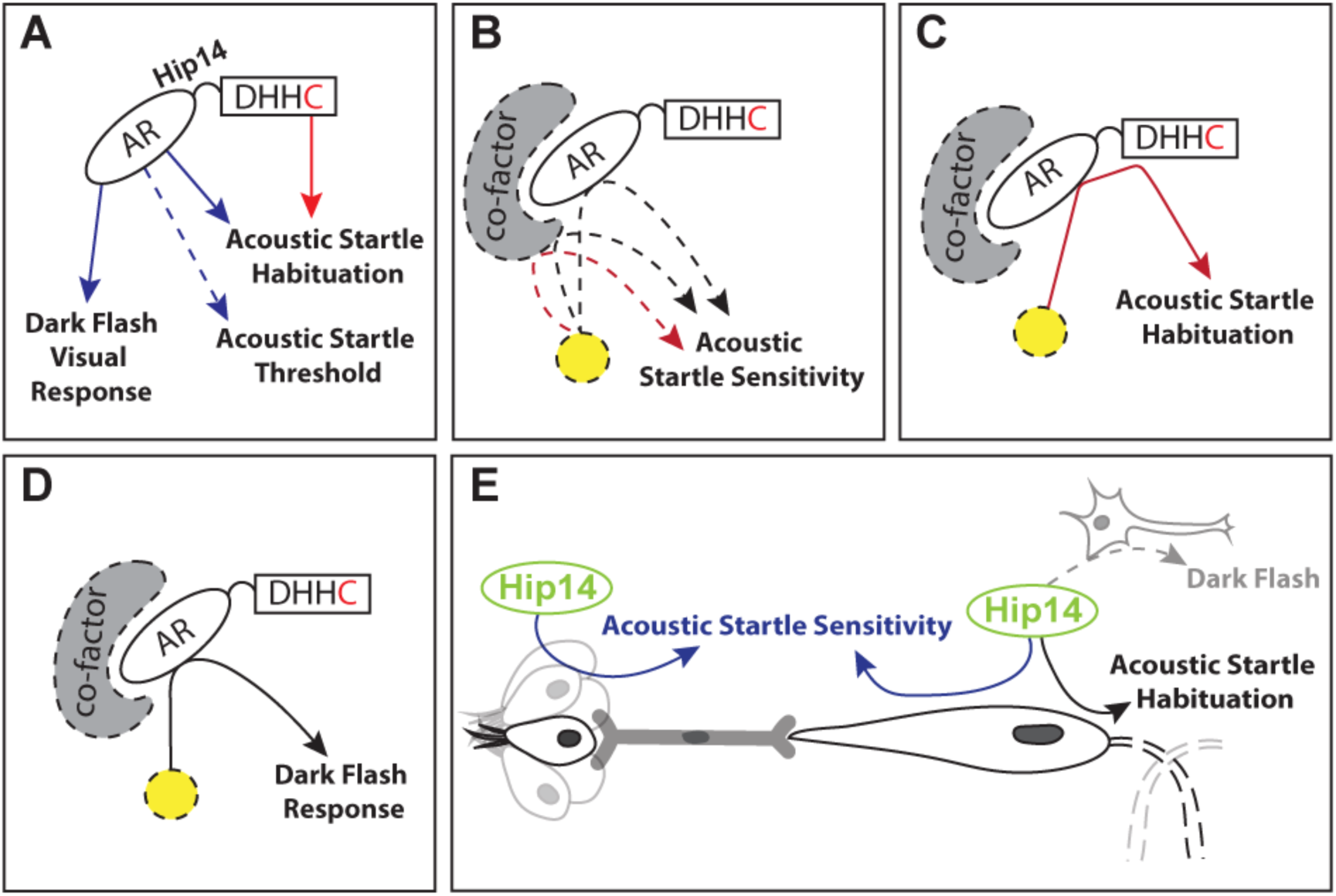
Model. (**A**) DHHC domain is required for acoustic startle habituation. AR domains are required for acoustic startle habituation and visual behaviors, but only partially required for acoustic startle sensitivity. Blue lines indicate Hip14 functions that are potentially DHHC-independent. Red line indicates proposed palmitoylation-dependent roles. (**B-D**) Proposed mechanism of Hip14 in regulating (**B**) acoustic startle sensitivity (black lines indicate Hip14 functions that are potentially DHHC-independent. Red dashed line indicates proposed palmitoylation-dependent roles), (**C**) acoustic startle habituation, and (**D**) Visual Response (dark flash). Even if the DHHC domain is dispensable, it is possible that a distinct PAT interacts with Hip14, possibly via the ankyrin repeats to promote Hip14-DHHC-independent palmitoylation. (**E)** Graphic display indicates where we propose Hip14 functions within the acoustic startle circuit. Hip14 can regulate acoustic startle sensitivity within both sensory hair cells and neurons.

Prior work demonstrates roles for Hip14 that are potentially independent of substrate palmitoylation (Yang and Cynader, 2011). These functions include cell cycle regulation (Hemsley and Grierson, 2011), golgiphagy (Nthiga et al., 2021), and JNK activation (Harada et al., 2003; Yang and Cynader, 2011; Yang et al., 2013; Chen et al., 2020). Other DHHC-independent roles still rely on protein palmitoylation, though not through the Hip14 DHHC (Goytain et al., 2008). We propose that, in the context of regulating acoustic sensory threshold behaviors, Hip14 could perform a palmitoylation-independent role and/or could interact with other PATs via the AR domain (**Fig.6B**). Several PATs are known to form dimers, such as zDHHC2 and zDHHC3 (Fang et al., 2006; Lai and Linder, 2013). zDHHC13 also potentially interacts with other PATs, although the specific interactor has not been specified (Petropavlovskiy et al., 2026). Beyond broadly identifying ankyrin repeat domains as important for behavioral regulation, we provide *in vivo* behavioral evidence for the importance of previously identified critical residues (N90/W118) (**Fig.5E-G**, **Fig.6B-D**).

### Hip14 is expressed and functional in mechanosensory hair cells

Hip14 mutants showed upregulated activity throughout the brain of the larval zebrafish, including within the lateral dendrites of the M-cells and spiral fiber neurons (Nelson 2023, 2020). Synaptic depression at the M-cell lateral dendrite (the site of inputs from the VIIIth nerve afferents) accompanies habituation learning, but in *hip14* mutants, activity is not reduced after repeated acoustic stimulation. These data are consistent with a model in which hyperexcitability upstream of the M-cells is responsible for reduced acoustic startle thresholds and habituation. Here, by examining endogenous Hip14 expression and using a transgenic rescue approach, we uncover that Hip14 is expressed both in neurons and in mechanosensory hair cells upstream of the M-cell. We find that, as expected, rescue of habituation requires re-expression of Hip14 in neurons. Expression in neurons is also sufficient to restore normal sensitivity to acoustic stimuli at baseline, but expression just in hair cells also partially restores sensitivity. These results are consistent with a model in which upregulated activity upstream of the M-cell renders larvae hyperresponsive to acoustic stimuli. Restoring Hip14 within neurons or mechanosensory hair cells can restore this upregulated activity to enable animals to gate incoming acoustic stimuli (**Fig.6E**).

### Hip14 is expressed in the Golgi in multiple cell types of the acoustic startle circuit

Our understanding of the underlying mechanisms that promote PAT substrate specificity is limited. But PAT subcellular localization could play a key role. Different PATs localize to distinct subcellular compartments. For example, zDHHC 3, 7, 13, 17, 21, and 22 are localized to the *cis-*Golgi, whereas zDHHC 9,11, and 15 are found in the *trans-*Golgi and post-Golgi structures (Ohno et al., 2006; Greaves et al., 2008; Globa and Bamji, 2017; Philippe and Jenkins, 2019). Furthermore, synaptic localization of DHHC2 is especially important for its function: synaptic activity triggers de-palmitoylation of PSD-95 at dendritic spines, which then relies on plasma membrane-localized DHHC2 for rapid re-palmitoylation to maintain postsynaptic nanodomain and AMPAR clustering (Fukata et al., 2013).

Though the majority of studies found zDHHC17/Hip14 primarily localized to the Golgi and, to a lesser extent, cytoplasmic vesicles (Singaraja et al., 2002; Huang et al., 2004), *Drosophila* HIP14 (dHIP14) and the human Hip14 homolog were found to localize to synapses (*in vivo* and in culture, respectively), co-localizing with synaptic vesicle proteins (Stowers and Isacoff, 2007). Similarly, other experiments identified dHIP14 localizing to the synaptic dense neuropil of the CNS and neuromuscular junctions (Ohyama et al., 2007; Stowers and Isacoff, 2007). Moreover, inhibition of synaptic vesicle endocytosis promoted re-localization of dHip14 to the synaptic plasma membrane, consistent with a localization of Hip14 not just to synapses but to synaptic vesicles (Ohyama et al., 2007). These studies raised the possibility of a direct role of Hip14 in regulating presynaptic plasticity. However, here we found that at least within the larval zebrafish brain, Hip14 is predominantly localized to the Golgi and not to synapses in both neurons and mechanosensory hair cells (**Fig.2**). These data contravene our hypothesis that local Hip14 function at synapses directly regulates neurotransmission relevant to sensory thresholds and habituation. Rather our findings are consistent with interaction of Hip14 with relevant substrates at the Golgi, followed by possible trafficking of protein interactors to synapses.

### Hip14 regulation of visual responses

Hip14 regulates acoustic sensory thresholds and habituation, but whether it also regulates processing in other sensory domains in vertebrates is largely unknown. To further explore Hip14 structure-function and circuit-level function, we assessed responses to dark flash (visual) in addition to acoustic stimuli (Burgess and Granato, 2007b). The O-bend response to dark flash stimuli is dependent on detection of visual stimuli at the level of the retina (Fernandes et al., 2012) and subsequent activity within the pallium, tectum, and cerebellum (Marquez-Legorreta et al., 2022), and ventromedial reticulospinal neurons required for turning responses (Orger et al., 2008; Huang et al., 2013). The initial screen identified visual habituation deficits in *p174* homozygotes (Wolman et al. 2015). However, our results show that *hip14^p174^* mutants exhibit deficits at the level of initial responsiveness: *hip14* mutants perform dark flash responses with strongly reduced frequency. In the context of this distinct behavioral deficit, Hip14 AR domains are required while the DHHC domain is dispensable (**Fig.4D & Fig.5D,G**). In the case of acoustic startle, our results demonstrate that Hip14 can function at the level of neurons and also peripheral sensory cells to control behaviors. In the case of visual behavior, we demonstrated that Hip14 plays a role in regulating responses via a function in neurons. However, given that our rescue was only partial, Hip14 could also be involved at the level of stimulus detection within the retina, synaptic transmission within photoreceptors, or other retinal cells as in *Drosophila* (Stowers and Isacoff, 2007).

Overall, our work demonstrates that establishment of acoustic startle thresholds and habituation learning, although often modulated together, rely on different genetic mechanisms and cellular processes. Hip14 regulation of acoustic sensitivity, acoustic habituation, and dark flash visual responses exhibit distinct requirements for individual protein domains, suggesting multiple parallel molecular functions for this key enzyme. Finally, our data highlight DHHC-independent functions *in vivo*. Future work will establish whether DHHC-independent functions are palmitoylation-independent or involve engagement of complexes including other PATs.

### Limitations

Our work primarily uses behavior as a readout for molecular function. We do not examine *in vivo* protein-protein interactions, although we speculate as to how they may be disrupted by our manipulations. Rescue transgenes use cell-type-specific promoters not expected to recapitulate the timing or levels of endogenous expression but rather designed to understand where and how Hip14 is sufficient to regulate behavior. Similar structure-function manipulations have been published in *in vitro* studies, and our ability to test a variety of behavioral assays establishes each construct’s functionality. Although the perinuclear localization of *hip14* in M-cells is convincing for Golgi localization **(Fig2.A-D)**, the *ubi-tdTomato-GM130* line failed to reach M-cells (**Fig.2CD**). Furthermore, although we did not observe synaptically localized Hip14, this does not eliminate the possibility that Hip14 subcellular localization is regulated by activity or other factors. It is also possible that synaptic Hip14 is present at a low level, below the detection threshold. Finally, we cannot distinguish between DHHC-independence and palmitoylation-independence. PATs can form complexes with one another, and one exciting possibility is that Hip14 promotes behavior independent of its DHHC domain by complexing with other PATs. No matter how Hip14 functions independent of its DHHC domain, our work provides evidence that the DHHC domain is strictly required for certain behaviors (habituation) and dispensable for others, highlighting the molecular versatility of this enzyme.

## Supplemental Figures

**Supplement Figure S1:**
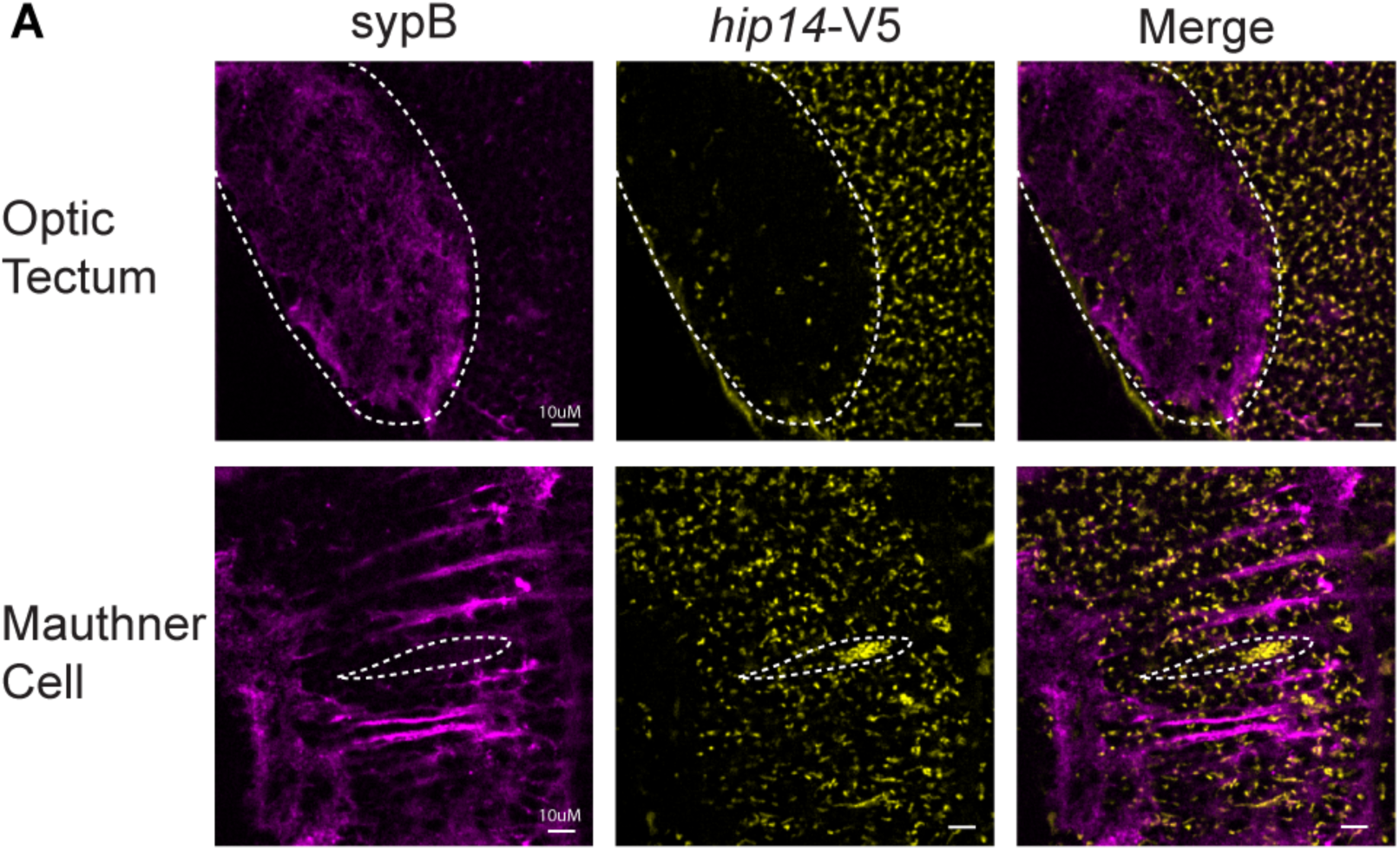
Hip14 is not presynaptic in optic tectum or hindbrain neuropil. **(A)** Hip14-V5 (yellow) does not colocalize with Synaptobrevin (SypB, in magenta) in the neuropil of optic tectum or hindbrain.

## Acknowledgments

We thank Dr. Katie Kindt for sharing the *p5e:myo6b* plasmid and the *myo6b:GCaMP6s:CAAX* line. We thank Dr. Bruce Appel lab for sharing the *NeuroD-sypB-mScarlett* plasmids. We thank Dr. Lila Solnica-Kretzel and Dr. Diane Sepich for sharing the GM130 plasmid. We are grateful to Dr. Koichi Kawakami and Dr. Jonathan Raper for sharing the *gffdmc130a* and *uas:gap43-citrine* lines. We would like to thank Teresa Nicolson for providing the anti-Vglut3 antibody. We thank the University of Colorado Anschutz Medical Campus Zebrafish Facility. We thank Dr. Victor A. Ruthig for microscopy support. This work is supported by funding from the NIH/NINDS Pathway to Independence Award (R00NS111736 to JCN), NIH/NIGMS Maximizing Investigators’ Research Award (MIRA R35GM159847 to JCN), NIH/ORIP R24OD037632 to CM and AB, and the Children’s Hospital Colorado Foundation to C.M. (The Helen and Arthur E. Johnson Chair For The Cardiac Research Director). The authors used ChatGPT to assist in writing code for statistical analyses presented in this manuscript. The authors tested and validated all code and accept full responsibility for all data and findings reported.

## Author Contribution

Designed research: Yuzhu Cheng, Jessica Nelson

Performed research: Yuzhu Cheng, Kade Fink, Nicholas Santistevan, Jessica Nelson Contributed unpublished reagents/analytic tools: Seth Jacobson, Alexa Burger, Christian Mosimann

Analyzed data: Yuzhu Cheng

Wrote the first draft of the paper: Yuzhu Cheng

Edited the paper: Yuzhu Cheng, Nicholas Santistevan, Kade Fink, Seth Jacobson, Christian Mosimann, Alexa Burger, Jessica Nelson

---

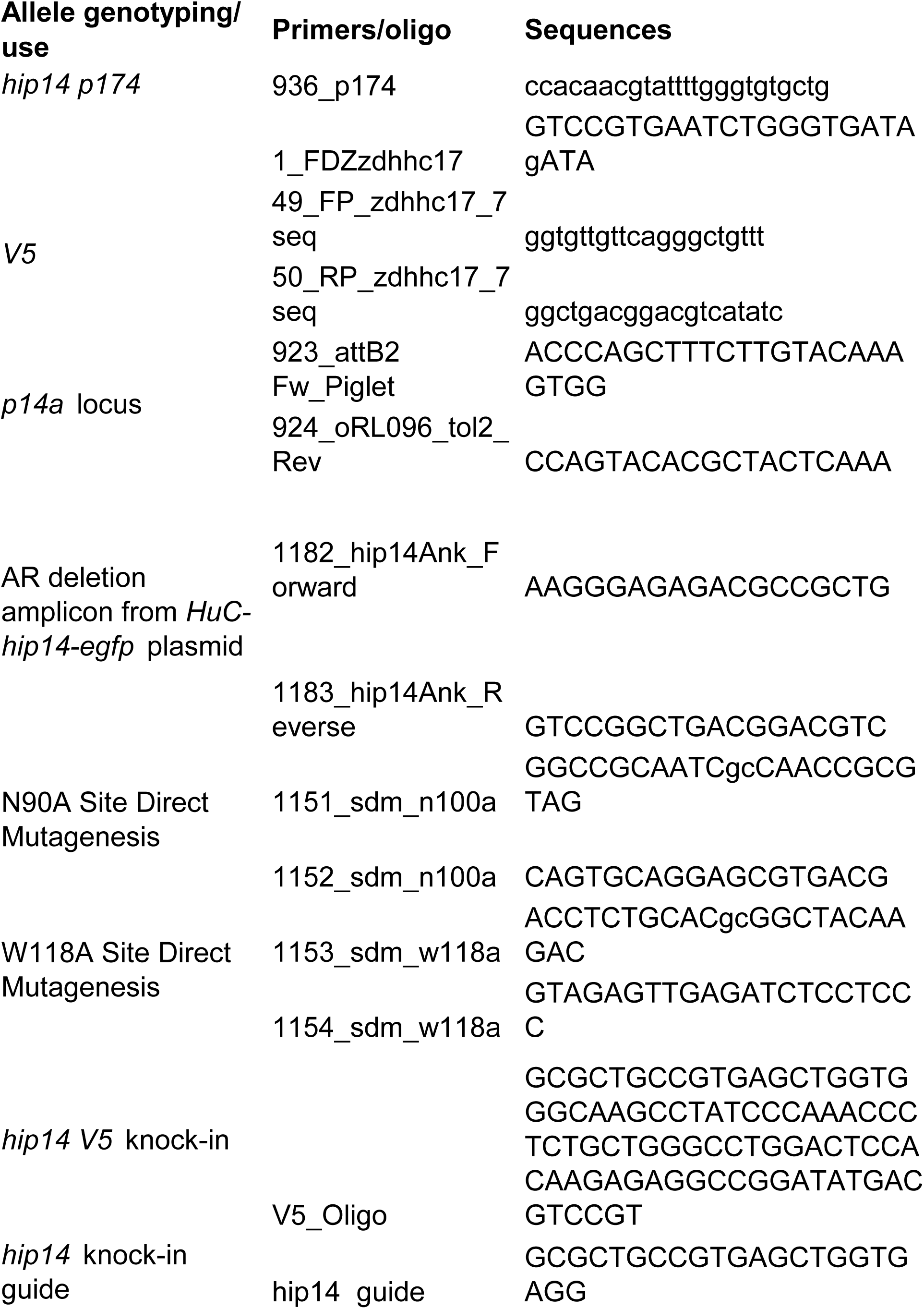
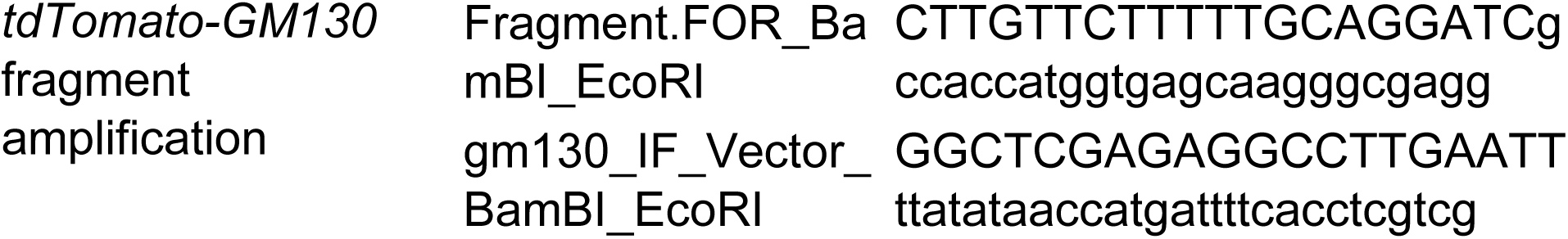

